# Genotypic and allelic distribution of the ACE, ACTN3, and IL6 genes in athletes of Artistic Gymnastics and Swimming in the department of Norte de Santander, Colombia

**DOI:** 10.1101/2022.06.07.495182

**Authors:** González Gerardo David, Correa William, Ruiz Casas Jairo

## Abstract

The purpose of the study is to report the allelic and genotypic frequencies of ACTN3, ACE and IL6 genes in Colombian high performance athletes in the sports of artistic gymnastics and swimming, to observe if there is any relationship between athlete status, sport and the polymorphisms found. The study included twenty-one gymnasts and twenty-one swimmers. The ACE and IL6 genes were genotyped by PCR and polyacrylamide electrophoresis, the ACTN3 gene was genotyped by PCR-RFLP. In the ACE gene, a greater number of genotypes with the D allele was observed in both sports, for the ACTN3 gene a predominance of the RX genotype was observed in both gymnastics and swimming, in the IL6 gene the most predominant characteristic is the total absence of the CC genotype. For the ACE gene, the highest allelic frequency is that of the D allele in both sports, the IL6 gene shows a high presence of the G allele, for the ACTN3 gene, in gymnastics the highest frequency of carriers is observed in the R allele and for the sport of swimming the majority is carrying the X allele. Significant differences were observed when comparing the genotypic and allelic frequencies between the population of gymnasts and swimmers only in the ACTN3 gene, the ACE gene and IL6 did not present significant differences between the populations. In conclusion, the genotypes presented by the athletes correspond to the types of efforts of their sport, gymnastics with predominance in strength and power and swimming with an orientation towards both phenotypes, showing a pleiotropic effect of the ACE and ACTN3 genes.

## INTRODUCTION

The advances in molecular biology that have been experienced in the last twenty years have allowed the development of knowledge in different areas, among these are sports sciences, which have benefited from advances in gene sequencing and their association with These physical activity, revealing databases that relate different genotypes with better athletic performance in different activities in which alactic, lactic anaerobic or aerobic resistance metabolisms predominate. Within these studies, artistic gymnastics and swimming have associated different genes with optimal performance in these sports, within them we have the ACE gene, which has been extensively studied in different sports, including artistic gymnastics (Morucci 2014) and have registered their relationships with the sport of high achievement, this gene presents a polymorphism (rs 1799752) represented in two alleles, D and I, the first of them has been related to good performances in strength activities, while the allele I has been associated with aerobic activities. In gymnastics, we have the relationship made by Fernandes Filho, Joao A.F. 2015, who studied the genetic profiles of Brazilian and Japanese athletes with this gene and its relationship with high-performance gymnastics. Another of the gene polymorphisms related to physical activity, which has been extensively studied, is the rs1815739 of the ACTN3 gene, known for its relationship with good results in power activities, this gene has a stop codon that prevents it from being completely transcribed, the protein alpha actinin 3, generating two alleles, the R which is associated with protein production and good performance in strength activities and the X allele, which represents the stop codon and the inactive protein, to which associated with good performance in aerobic activities, within the studies carried out in artistic gymnastics and the ACTN3 gene, we found that of Massida et al. 2009, in which they associate the ACTN3 R577X polymorphism with good performances in male and female elite gymnastics.

The −174 G/C polymorphism (rs1800795) of the IL6 gene is related to immune functions and in the field of sports with muscle repair and hypertrophy, there is an association between the C allele of this polymorphism and high levels of creatine kinase after physical exercise, which is an indicator of muscle damage. On the contrary, it happens with the possessors of the G allele, Yamin et. al. We hypothesize that the R alleles of the ACTN3 gene, D of the ACE gene and G of the IL6 gene are found in higher allelic frequency in Olympic gymnastics and swimming athletes, since they provide competitive advantages in these sports.

## MATERIALS AND METHODS

The study population consisted of twenty-one gymnasts, among which we highlight the Colombian Selection of Men’s Artistic Gymnastics (GAM) Seniors (5 members) between the ages of 18 and 27, third by teams in the 2021 Pan American Games in Rio de Janeiro (champion in high bar, runner-up in parallel bars and pommel horse, runner-up in high bar), the Colombia Junior Men’s Artistics Gymnastics Team (5 members) between the ages of 14 and 17, Pan American Runners-up (by teams, Pan American champions in high bar, third on floor, bar and all around, and parallel runner-up) runners-up Junior Games Guadalajara July 2021 and Champions of the South American Sports Games Rosario April 2022. Five gymnasts from the Junior category between 16 and 17 years old, eleven Junior MAG Gymnasts and Seniors, all of them national medalists and twenty-one swimmers from the Norte de Santander Swimming League, aged between 14 and 17, all of them within the 8 first places in the FECNA National Ranking (Colombian Swimming Federation) in their respective categories, some of them national gold medalists in the National League tournaments endorsed by this Federation.

The project adhered to the protocols required in studies carried out in humans, by the Helsinki protocols and according to resolution 008430/1993 of the Colombian Ministry of Health. Prior to taking the sample, each athlete signed an informed consent, where they agreed to participate in this study.

DNA from buccal scraping samples was extracted by implementing the protocol used by Quinque et al (2006). After DNA extraction, the extracted samples were quantified using the Thermo Scientific NanoDrop2000/2000c spectrophotometer. Depending on the concentration in ng/μl found in each sample, the concentration was adjusted to 20 ng/μl. The genotyping of each of the loci was performed by amplification by the PCR technique, from the DNA located in autosomal chromosomes, with the following primers: Gene ACE Forward 5’CTGGAGACCACTCCCATCC1TTCT 3’, Reverse 5’ GATTGGCCATCACATTCGTCAGAT 3’, Gen ACTN3 Forward 5’-CTGTTGCCTGTGGTAAGTGGG-3’, Reverse 5’-TGGTCACAGTATGC AGGAGGG-3’ Gen IL6 Forward 5’-ATAAATCTTTGTTGGAGGGTGAGG 3’, Reverse (allele C)5’ATGACGACCTAAGCTTTACTTTTCCCTACTG-3,Reverse(alleleG)5’-GCACTTTTCCCCCTAGTTGTGTCTTACG-3

The following amplification protocol was used for all genes, which consists of the following cycles, 95.0 °C, 4:00 min, 35 cycles, 94.0 °C, for 1:00 min, 58.0 °C for 1:00 min, 72.0 °C for 1:00 min, followed by a final elongation of 72.0 °C, for 10:00 min. The PCR reaction will be performed at a volume of 25μl, which will contain 3 mM MgCl2, 0.1 mM primers, 0.1 mM dNTPs, 1X buffer, 1X Taq polymerase. The amplified fragments of the ACTN3 gene were enzymatically digested with their corresponding restriction enzyme, Dde1. Subsequently, electrophoresis was performed in 8% polyacrylamide gels, in TBE1X of the products obtained in the PCR. These electrophoresis will be run at 130 volts, and will be visualized on gels stained with silver nitrate.

### Statistical analysis

The genotypic frequencies of the athletes will be evaluated in compatibility with the Hardy Weinberg equilibrium, the genotypic distribution and the allelic frequencies among the group of artistic gymnastics and swimming athletes will be calculated and compared between them, their significance tested by *X^2^* (p <0.05). The normality of the data will be verified with the Kolmogorov-Smirnov test. For the above calculations, the STATISTICA 8.0 software (StatSoft Inc 2007) will be used.

## RESULTS

The genotyping of the population was carried out according to the established methodology. In table 1, it is observed in the ACE gene, a greater number of genotypes with the D allele in both sports, despite not presenting significant differences when comparing these genotypes between the sports of gymnastics and swimming, P=0.2397 (table 3), with the difference that the DD genotype predominates in gymnastics and the DI genotype in swimming, in addition, no individual of the two sampled populations presents genotype II. For the ACTN3 gene, a predominance of the RX genotype is observed in both gymnastics and swimming, followed by the RR genotype for gymnastics and XX in swimming. When observing the IL6 gene, the most predominant characteristic is the total absence of the CC genotype and the majority of individuals with the GG genotype in both sports.

**Table 1.** Genotypes presented in the athlete populations of the Colombian artistic gymnastics team and the swimming team of the department of Norte de Santander

| GENOTYPES |  |  |  |  |  |  |  |  |  |
| --- | --- | --- | --- | --- | --- | --- | --- | --- | --- |
|  | ACE |  |  | ACTN3 |  |  | IL6 |  |  |
|  | DD | II | ID | RR | XX | RX | GG | CC | GC |
| **Gymnastics** | 12 | 0 | 9 | 4 | 2 | 15 | 16 | 0 | 5 |
| **swimming** | 8 | 0 | 13 | 3 | 8 | 10 | 15 | 0 | 6 |

**Table 2.** Allelic frequencies presented in the populations of athletes from the Colombia and North Artistic Gymnastics teams and the Departmental Swimming team from Norte de Santander.

| ALLELIC FREQUENCIES |  |  |  |  |  |  |
| --- | --- | --- | --- | --- | --- | --- |
|  | ACE |  | ACTN3 |  | IL6 |  |
|  | D | I | R | X | G | C |
| **Gymnastics** | 0,786 | 0,214 | 0,548 | 0,452 | 0,881 | 0,119 |
| **Swimming** | 0,690 | 0,310 | 0,381 | 0,536 | 0,857 | 0,143 |

**Table 3.** Comparison between genotypes and allelic frequencies between the sports of gymnastics and swimming

|  | Gymnastic vs Swimming |  |  |
| --- | --- | --- | --- |
|  | ACE | ACTN3 | IL6 |
| **P genotype** | 0,2397 | 0,0141 | 0,8474 |
| **P alleles** | 0,2427 | 0,0389 | 0,8255 |

Regarding the allelic frequencies (Table 2) for the ACE gene, the highest frequency is that of the D allele in both sports, the same occurs with the IL6 gene and the allele G, showing a fairly high prevalence with respect to the allele. Regarding the ACTN3 gene, in gymnastics the highest frequency of carriers is observed in the R allele and for the sport of swimming, most are carrying the X allele.

When comparing the genotypes and allelic frequencies presented in the population of gymnastics athletes against the genotypes and allelic frequencies of swimming athletes, significant differences are observed between the genotypes of the ACTN3 gene and the allelic frequencies of this same gene. No significant differences were observed in the ACE and IL6 genes between the two populations.

## DISCUSSION

This is the first study conducted in high-performance gymnastics and swimming athletes in Colombia, with three genes (ACE, ACTN3 and IL6), which have been extensively investigated in the field of sport genetics. In the artistic gymnastics group are the Colombian national team athletes, the most representative with international achievements between 2015 and 2022; This sport is a competitive art sport that is characterized by a predominance of central, neuromuscular control, with high motor potential, mediated by the conditional directions of force, determining factors for its sporting performance, especially power and maximum force, all of these under anaerobic conditions, ranging from alactic anaerobic (phosphogens) to lactacid anaerobic (anaerobic glycolysis) given the level of effort in which their competition tests result, ranging from the “Vault Jump” with an approximate duration of 10 seconds, until the “Floor exercises” with an approximate duration of 90 seconds, which forces the constant stimuli that the athlete receives throughout his sports career to generate epigenetic factors that select genes oriented to the development of these specific capacities mentioned above; swimming, on the other hand, is a cyclical sport, with an aerobic base, mediated by conditional resistance capacities, in which the determining directions for high sports performance are based on mixed conditions (aerobic anaerobic) that combine both anaerobic conditions (lactic and alactic, in competition tests ranging from 50m to 200m), to tests with aerobic predominance ranging from VO2MAX (aerobic power), Maximum Lactate Stable States (Maxlass or MLSS), to aerobic conditions long-term, that is, from 400 meters in confined water (swimming pool) and up to 10,000 meters in open water (from aerobic glycolysis to mitochondrial pathways of lipolysis, gluconeogenesis, among others), It is very important to highlight in swimming that the basis of training throughout a macrocycle is predominantly aerobic, especially in the ages of sports development (12 to 16 years of age), which forces the athlete who projects high performance, to receive daily stimuli, throughout their sports career, that select genes oriented to the development of this combination of mixed capacities with an important aerobic component. In a nutshell, while gymnastics is a central control sport dominated by strength conditionals, swimming is a peripheral control sport that seeks metabolic adaptations dominated by mixed resistance conditionals, so that, in each sport, they will mediate specific epigenetic factors that must be activated, both polymorphisms and alleles, aimed at a much more efficient sports selection on their way to high performance.

It is important to note that, whatever the sport that is practiced, if you want to reach high performance, the athlete is subjected to constant metabolic and physical stress given the frequency of training sessions in a microcycle, (more than 7 per week) that forces the athlete to generate constant and rapid recovery processes in order to be ready to bear the load of the next training session, therefore, both homeostasis and super-compensation processes require a special sports selection in subjects who efficiently adapt to these work rhythms, becoming more efficient in these recovery genetic actions on the way to high performance.

The analysis of the groups allows us to observe how, thanks to sports selection, exercised in athletes, most of the genotypes present in gymnastics are related to those previously reported in the literature (Cieszczyk et. al. 2011. Eynon et. al. 2013, Ma et. al. 2013), as beneficial for strength activities. The genotypes DD of the ACE gene, RX of the ACTN3 gene and GG of the IL6 gene are the ones with the highest appearance in gymnastic athletes (DD=12; RX=15; GG=16). In addition, these genotypes have already been reported in previous publications (Joao et al 2015. Myosotis et al 2009. Gabriele 2014) as those with the highest appearance in gymnastic athletes. Regarding the allelic frequencies, as with the genotypes, the alleles related to force activities in previous publications (Eider et. al. 2013, Ruiz et. al. 2010) such as the D allele of the ACE gene, the R allele of the ACTN3 gene and the G allele of the IL6 gene, have a greater presence in the athletes of the Colombian gymnastics team (D=0.78; R=0.54; G=0.88). All of the above may be due to the sport’s orientation towards anaerobic lactic and alactic metabolisms, related to power activities, maximum strength and resistance to force (anaerobic?), which are predominant in the sport of gymnastics.

In relation to swimming athletes, it is observed that in the genotypes, related to the ACTN3 and ACE genes, the greatest presence is of the heterozygotes, DI for the ACE gene and RX for the ACTN3 gene (DI=13; RX=10), perhaps related to the aerobic component that this sport has and alleles I of the ACE gene and X of the ACTN3 gene, reported in previous publications (Chiu et. al. 2011. Wang et. al. 2013), the predominant genotype of the IL6 gene, was the GG (GG=15) reported as beneficial for strength activities, which can improve sports performance in sprint moments of swimming competitions, together with the RX and DI genotypes of the ACTN3 and ACE genes respectively. When analyzing the allelic frequencies in the swimming population, a greater appearance of allele frequencies oriented to good aerobic performance is observed, such as the X allele of the ACTN3 gene, which is present in the majority in these athletes, and also a greater appearance of alleles that provide good performance in strength activities such as the D of the ACE gene and the G of the IL6 gene, which would provide suitable achievements in the parts of the competition where this quality is needed, such as the starts and the final sprints, in addition to the work of resistance to force, constant throughout the competition path.

The foregoing shows a possible relationship between the ACE and ACTN3 genes and pleiotropy, relating good performances in strength and resistance work, with the heterozygotes of these genes, which provide orientation in different phenotypes such as strength allele D gene ACE and allele R of the ACTN3 gene and aerobic resistance, allele I of the ACE gene and allele X of the ACTN3 gene, which can be seen in Table 1, where the majority of individuals present the RX genotype of the ACTN3 gene in both sports and the DI genotype in swimming, which is a sport that needs both characteristics in greater proportion than gymnastics.

Regarding the muscle recovery capacity related to the G allele of the IL6 gene, we observed in Table 1 a greater appearance of the GG genotype, which has been reported (Huuskonen et. al. 2009) as beneficial for muscle recovery in athletes, and there is no sampled athlete carrying the CC genotype, which is consistent when relating high performance with a good recovery capacity between series and daily training densities in mesocycles, which these athletes must endure.

When making a comparison of the genotypes of the three genes by means of the *X*^2^ test between the population of gymnasts and swimmers, we observed significant differences only in the ACTN3 gene (p=0.0141), the same occurring when comparing the allelic frequencies, where significant differences are only seen in the ACTN3 gene (p=0.0389), which may be due to the greater influence of alpha actinin 3 in strength activities, which is the predominant characteristic in gymnastics, and what would differentiate these two sports for the most part. A possible cause of not presenting significant differences in the ACE and IL6 genes could be the size of the sample.

In conclusion, the genotypes presented by the athletes correspond to the types of efforts of their sport, gymnastics with a predominance of strength and power and swimming with an orientation towards both phenotypes, showing a pleiotropic effect of the ACE and ACTN3 genes, together with the As has been reported, the IL6 gene provides, in addition to better performance in strength, a good post-training recovery at the skeletal muscle level, which suggests that genetic analysis is an excellent input for coaches and the development of individualized training plans. as well as for sports selection at an early age, aimed at high performance.

## ACKNOWLEDGMENTS

Norte de Santander Swimming League for their financial contributions.

Clubes Deportivos Cazadores Vida en el Agua y Focas who contributed their swimmers for the sample Liga Nortesantandereana de Gimnasia, for the contribution of its gymnasts

Colombian Federation of Gymnastics for the contribution of their teams

INDENORTE and the government of Norte de Santander for their economic contributions and support for this study

Parents of athletes for their financial support

## Notes

### Competing Interest Statement

The authors have declared no competing interest.

